# Evolution of the Cytochrome P450 Family 4 (CYP4) Subfamilies in Birds

**DOI:** 10.64898/2026.02.03.700598

**Authors:** Diksha Bhalla, Vera van Noort

## Abstract

Cytochrome P450 (CYP) genes form a large and functionally diverse superfamily of heme-thiolate monooxygenases that catalyse the oxidation of endogenous and exogenous compounds. Within this superfamily, CYP4 enzymes primarily act on fatty acids, substrates central to energy metabolism. Sustained flight and long-distance migration in birds rely on efficient use of fatty acids, making CYP4 a pertinent system for examining how lipid-associated functions may vary across avian lineages. Across Aves, species occupy diverse ecological niches and display substantial variation in metabolic demands. Here, we characterised CYP4 evolution across birds using 363 avian whole genomes, which cover over 90% of bird families, to investigate how gene variation relates to differences in ecological traits and metabolic strategies. Using this extensive genomic dataset, we identified 4 CYP4 subfamilies and analysed their patterns of sequence evolution. The analyses revealed conserved elements of the avian CYP4 repertoire alongside lineage-specific variation. Several amino-acid positions under positive selection were located within substrate recognition sites (SRS), regions that influence substrate binding and catalytic properties. Changes at these positions may reflect shifts in enzymatic function associated with differences in ecological or physiological traits among species. Overall, the results highlight SRS-associated variation within CYP4 that may reflect adaptive responses to environmental change. These findings advance understanding of how lipid-associated metabolic pathways have been shaped during avian diversification and provide insight into the evolutionary history of a CYP family linked to metabolic adaptation in birds.

## Introduction

Cytochrome P450 (CYP) enzymes constitute a large and diverse superfamily of heme-thiolate monooxygenases involved in the oxidation of endogenous substrates and environmental toxicants (Nelson and Nebert 2018). CYP enzymes are found across all domains of life and play central roles in metabolism, detoxification, and physiological regulation. In vertebrates, CYP genes have diversified into multiple families and subfamilies through gene duplication and loss, giving rise to lineage-specific repertoires with distinct metabolic functions. CYP proteins exhibit sequence diversity across families and species, reflecting functional specialisation within the superfamily. This diversification has occurred alongside conservation of key features required for heme-dependent monooxygenase activity across taxa (Werck-Reichhart and Feyereisen 2000; Guengerich 2015). CYP families and subfamilies are defined primarily by amino-acid sequence identity according to established nomenclature guidelines: sequences sharing >40% identity are assigned to the same family, while those with >55% identity are grouped into the same subfamily (Nelson et al. 1993; Nelson et al. 1996; Nelson 2005).

Among vertebrate CYPs, families CYP1–3 participate extensively in xenobiotic biotransformation while also acting on a range of endogenous molecules. CYP4 enzymes catalyse the ω-hydroxylation of fatty acids and lipid-derived signaling compounds (Ioannides 2008). Among the 72 known CYP4 subfamilies, 7 subfamilies have been identified in vertebrates: 4A, 4B, 4F, 4T, 4V, 4X and 4Z. Vertebrate CYP coding transcripts are approximately 1500 base pairs (bp) long and CYP4 genes typically comprise eleven to thirteen exons (Nelson 2003; Hardwick 2008). Studies in mammals have shown that CYP4A, CYP4B, and CYP4F subfamilies metabolise short-, medium-, and long-chain fatty acids, contributing to processes involved in fatty-acid turnover and the regulation of lipid homeostasis (Johnson et al. 2015). Although the roles of certain subfamilies, such as CYP4V, remain less clearly defined, the involvement of CYP4 enzymes in fatty-acid modification suggests potential relevance in organisms with substantial lipid-based metabolic demands.

Lipid-based energy metabolism is central to avian physiology, particularly in species that perform sustained flight or long-distance migration. Many migratory birds accumulate large fat reserves before sustained migratory movements and rely on the rapid mobilization and oxidation of fatty acids to support extended intervals of aerobic activity. These metabolic processes require coordinated regulation of lipid uptake, transport, and catabolism within oxidative skeletal muscle, followed by restoration of fuel stores between migratory flights. Studies suggest that fat provides up to 90% of the energy for migratory flight, while protein accounts for the remaining 10% (Jenni and Jenni-Eiermann 1998; Jenni-Eiermann et al. 2014; Araújo et al. 2019; McWilliams et al. 2021). The prominence of lipid utilization in birds has motivated extensive research on avian energy metabolism (McWilliams et al. 2004; Guglielmo et al. 2018; Eikenaar et al. 2022), yet the genetic and evolutionary components underlying fatty-acid processing pathways—particularly those involving CYP4 enzymes—remain poorly characterised. Given the known roles of CYP4 enzymes in mammalian fatty-acid oxidation, examining this family in birds may offer insight into its diversification in association with avian metabolic traits.

Comparative genomic analyses of avian CYP families have primarily focused on CYP1–3. Studies in chicken (*Gallus gallus*), zebra finch (*Taeniopygia guttata*), and turkey (*Meleagris gallopavo*) have described variation in CYP gene number and expression among lineages (Watanabe et al. 2013; Ren et al. 2019). In addition, a survey of the CYP2 family across 48 avian genomes identified heterogeneity in selective pressures and associations with ecological traits (Almeida et al. 2016). In contrast, the evolutionary history of the CYP4 family in birds remains largely uncharacterised, and how CYP4 genes have diversified across avian lineages remains unclear.

Here, we identified CYP4 genes across 363 avian whole genomes from 37 orders, including those from the Bird 10,000 Genomes (B10K) Project (Feng et al. 2020), which together represent over 90% of avian families, and analysed their phylogenetic patterns and adaptive evolution, providing a comprehensive framework for CYP4 diversification.

## Results

### Avian CYP4 subfamilies

The BLAST searches performed on 363 avian whole genomes identified a total of 1056 avian CYP4 sequences. Based on sequence similarity and domain composition, four CYP4 subfamilies were recovered: CYP4A, CYP4B, CYP4F and CYP4V (Fig 1).

**Fig 1.**
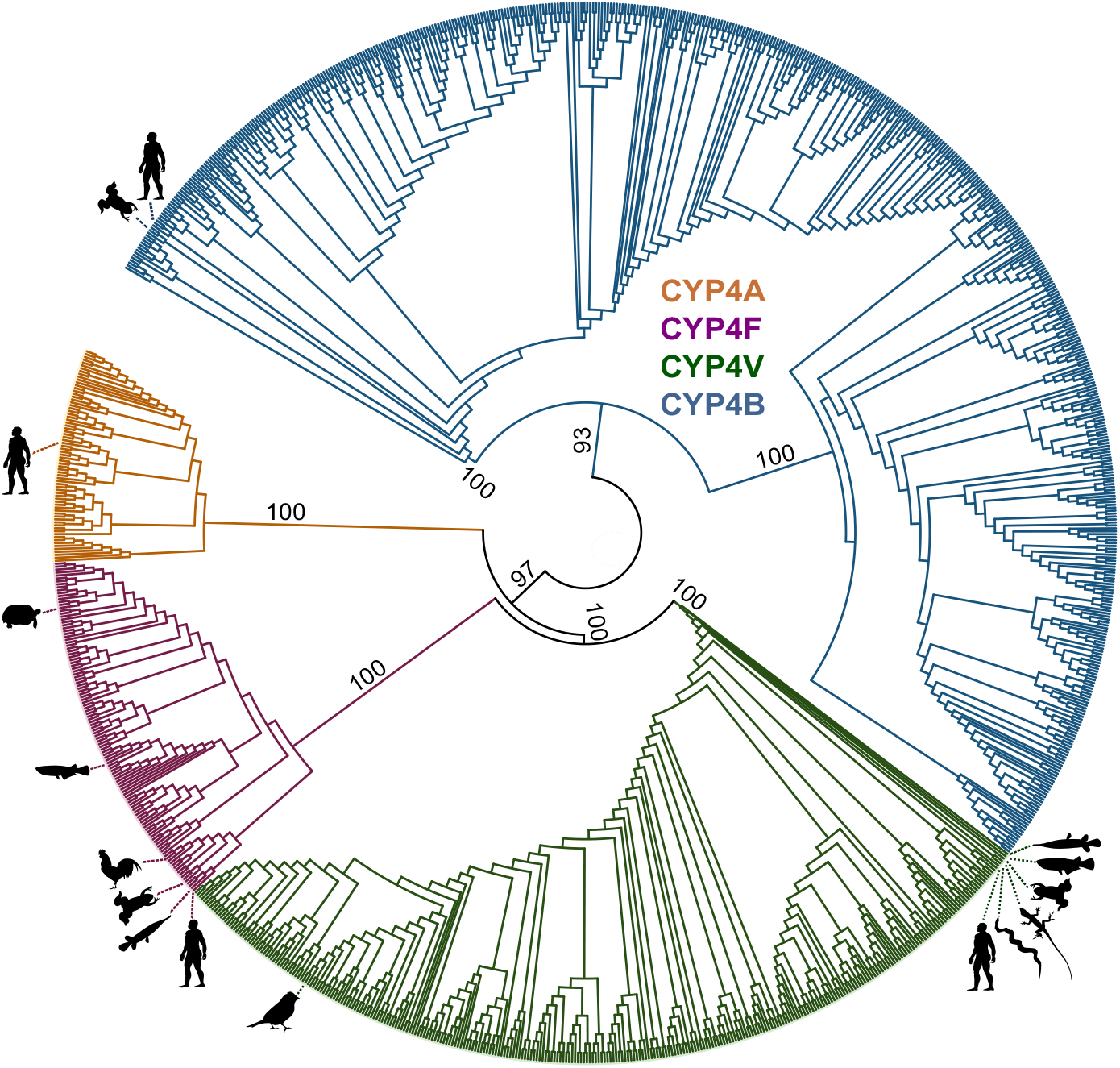
The Phylogenetic tree was built in IQTree (v2.3.6) using the ML method with 1000 bootstrap replicates. The tree represents the evolutionary history inferred from 1056 avian CYP4 nucleotide sequences from 363 avian genomes and 42 CYP4 nucleotide sequences of reference species from public databases. Bootstrap score is shown next to the branches.

These included 66 CYP4A genes identified in 65 species, 582 CYP4B genes in 340 species, 103 CYP4F genes in 93 species, and 305 CYP4V genes in 300 species.

Our analysis identified CYP4A genes in 65 avian species, including CYP4A22. However, this subfamily dataset did not meet the criteria described below for downstream codon-based analysis. While CYP4B, CYP4F, and CYP4V are broadly conserved across vertebrates,CYP4A has generally been characterised as a mammal-

specific subfamily in curated domain resources such as the NCBI Conserved Domain Database (CDD, entry cd20628). In chicken, CYP4A22 has previously been detected in liver transcriptome analyses and shown to be estrogen responsive (Ren et al. 2019), but the broader distribution and evolutionary context of CYP4A genes across avian geneomes has not been systematically examined.

### Evolution of CYP4 Subfamilies Among Birds

The ancestral reconstruction of CYP4 subfamilies performed in COUNT (Csűös 2010) suggested that the most recent common ancestor of extant birds must have harboured elements of CYP4A, CYP4B, CYP4F and CYP4V. Multiple CYP4 subfamilies are predicted to have been present at deeper ancestral nodes, with subsequent lineage-specific losses occurring across different avian clades. Among the CYP4 subfamilies, CYP4B and CYP4V show broader retention across the avian phylogeny compared with other subfamilies. Subfamily losses are distributed across both deep and terminal branches, and several species show evidence of subfamily loss at the tips. In some taxa, no CYP4 subfamilies were detected using the current criteria, which likely reflects incomplete chromosomal assemblies rather than true absence.

### Site-specific Signatures of Selection in Avian CYP4 Enzymes

To characterise patterns of molecular evolution across avian CYP4 genes, site models implemented in CODEML were used to test for heterogeneity in selective pressures among codon sites and to identify evidence for positive selection. Likelihood ratio tests (LRTs) were performed for standard model comparisons (M0 vs M3, M1a vs M2a, M7 vs M8, and M8 vs M8a), allowing discrimination between variation in selective constraint and support for codon classes evolving with ω > 1. Codons inferred to be under positive selection were identified using Bayes empirical Bayes (BEB) analysis, and Property Informed Models of Evolution (PRIME) were applied to examine whether selection at these sites was associated with changes or conservation of specific amino-acid properties (Table S2-S12). Where relevant, positively selected sites were examined in relation to predicted secondary structure and previously defined substrate recognition sites (SRS).

#### CYP4A4

Site-model analyses of CYP4A4 revealed significant heterogeneity in selective pressures among codon sites, with the discrete model M3 fitting the data significantly better than the one-ratio model M0 (Table S2). However, comparisons explicitly testing for positive selection (M1a vs M2a and M8a vs M8) were not significant, and the estimated ω for the additional site class under M8 was equal to one. These results indicate variation in selective constraint across sites but do not support the presence of a distinct class of codons evolving under positive selection. Despite the lack of robust model-level support, BEB analyses identified a single codon (site 337) with high posterior probability under both M2a and M8 (PP > 0.99; Table S3), corresponding to <0.3% of sites in the alignment. This site is located within a predicted helical region but does not overlap a previously defined SRS. PRIME analysis indicated that substitutions at this position were associated with changes in chemical composition and charge, alongside conservation of properties related to secondary structure and residue volume. These findings suggest that selective pressures at this site may be constrained by local structural context, although evidence for broader adaptive evolution in CYP4A4 remains limited.

#### CYP4A5

In CYP4A5, all site-model comparisons consistently supported site-specific positive selection. The M3 model indicated substantial heterogeneity among sites (Table S4), and both M1a vs M2a and M7 vs M8 comparisons supported models allowing a positively selected site class. This signal was further confirmed by a significant M8a vs M8 comparison. Under M2a, approximately 5.9% of sites were assigned to the positively selected class (p_2_ ≈ 0.059), with an estimated ω_2_ exceeding 2. BEB analyses identified nine codons with high posterior support for positive selection (PP ≥ 0.95), distributed across the protein (Table S5). Several of these sites were located within predicted helices, including residues overlapping substrate recognition sites SRS1 (positions 106 and 112) and SRS6 (position 479). PRIME analyses detected limited property-level signals, with only one site showing significant conservation of secondary-structure-related properties under the applied thresholds. Overall, these results indicate that positive selection in CYP4A5 affects a modest proportion of sites, including residues within functionally relevant regions, while many substitutions remain physicochemically conservative. Given the presence of positively selected sites within both SRS1 and SRS6, the spatial distributuin of these residues was examined using homology-based structural models to assess their relative positioning within the protein (Fig 2).

**Fig 2.**
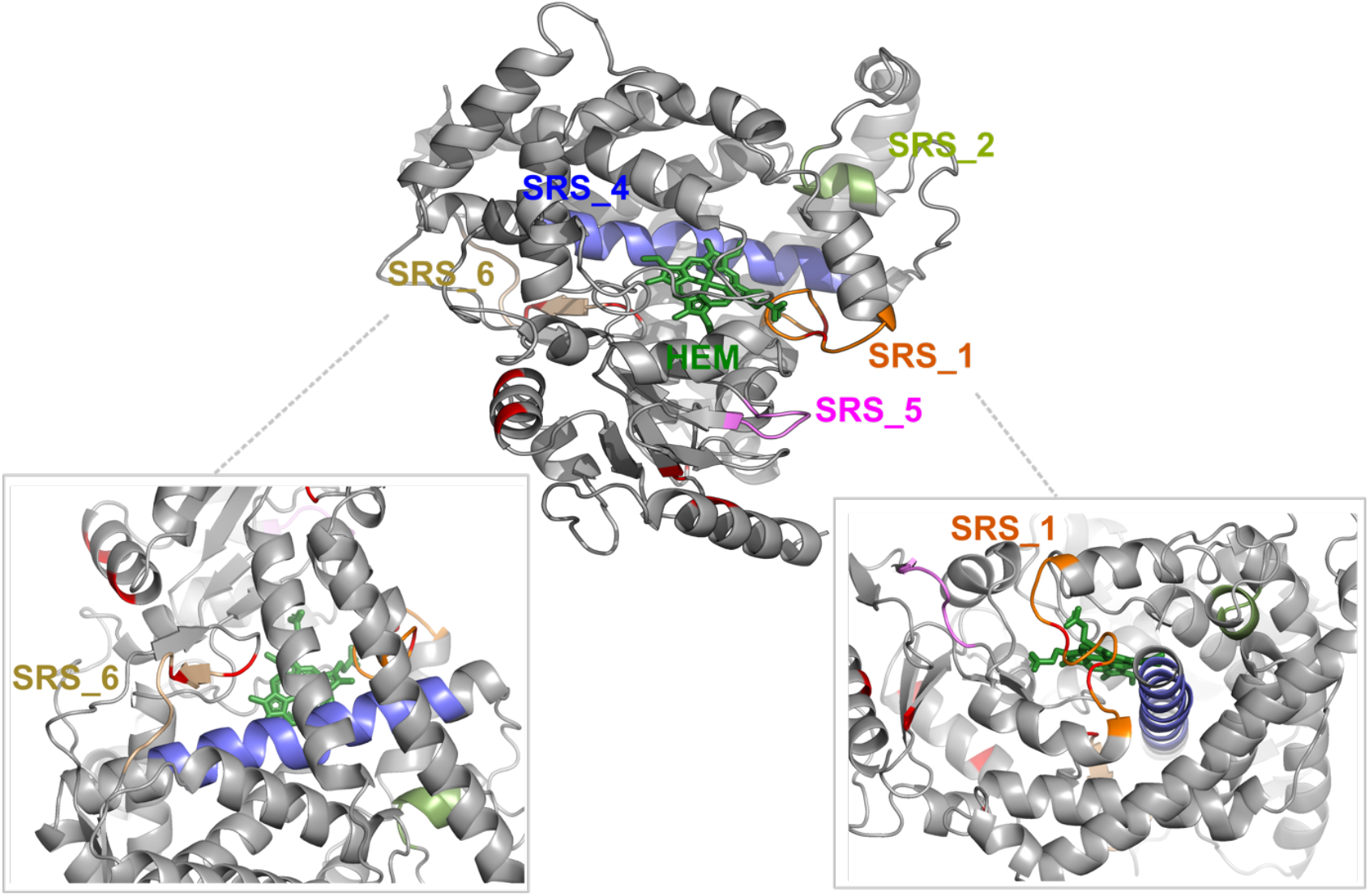
3D analysis of sites identified as being under positive selection in the avian CYP4A5. Positively selected sites identified by codeml site-models analyses (BEB, PP ≥ 0.95) are mapped onto a homology-based structural model of CYP4A5, shown in red. Residues overlapping previously defined substrate recognition sites (SRS) are hihglighted. SRS 1: orange, SRS 2: light green, SRS 4: blue, SRS 5: magenta and SRS 6: wheat.

#### CYP4B1

CYP4B1 exhibited strong and concordant evidence for site-specific positive selection. The M3 model significantly improved fit relative to M0, indicating pronounced heterogeneity in ω across codon sites (Table S6). Comparisons allowing for positive selection (M1a vs M2a and M7 vs M8) were highly significant, and the M8a vs M8 test further supported the presence of sites evolving with ω > 1. Under M2a, approximately 2.2% of sites were assigned to the positively selected class (p_2_ ≈ 0.022), with ω_2_ estimates exceeding 2. BEB analyses identified multiple codons with high posterior support for positive selection (PP ≥ 0.95), including sites 201, 205, 261, 318, 324, 339, 342, and 347 (Table S7). Several of these sites mapped to predicted helices, and at least one site (318) overlapped substrate recognition site 5 (SRS5). PRIME analyses provided site-specific characterisation of the physicochemical properties associated with substitutions at these positions. Site 201, located within a predicted helix, showed evidence for changes in amino-acid composition and residue volume, while properties related to secondary structure and bipolarity were conserved. Site 205, also located within a helix, was associated with changes in charge, with conservation of bipolarity. Site 261, not annotated to a specific secondary structure element, exhibited changes in amino-acid composition and secondary structure related properties, alongside conservation of charge and bipolarity. Site 318, which overlaps SRS5, showed changes in amino-acid composition, with conservation of bipolarity and charge-related properties. Site 324, lacking an explicit secondary-structure annotation, was associated with changes in composition, while bipolarity and residue volume were conserved. Site 339, also without a specific structural annotation, showed changes in composition, with conservation of bipolarity. Site 342, located within a predicted β-strand, exhibited changes in secondary structure related properties, while composition and bipolarity were conserved. Site 347, located within a predicted helix, showed changes in secondary structure related properties, with conservation of bipolarity, composition, and charge.

Taken together, these results indicate that positive selection in CYP4B1 affects defiend set of sites distributed across helices, strands, and substrate recognition region, with substitutions associated with site-specific shifts in physiochemical properties. Given the concentration of positively-selected sies within structured regions and within SRS5, their spatial distribution was examined using homology-based structural models to infer their relative position within the preotin (Fig 3).

**Fig 3.**
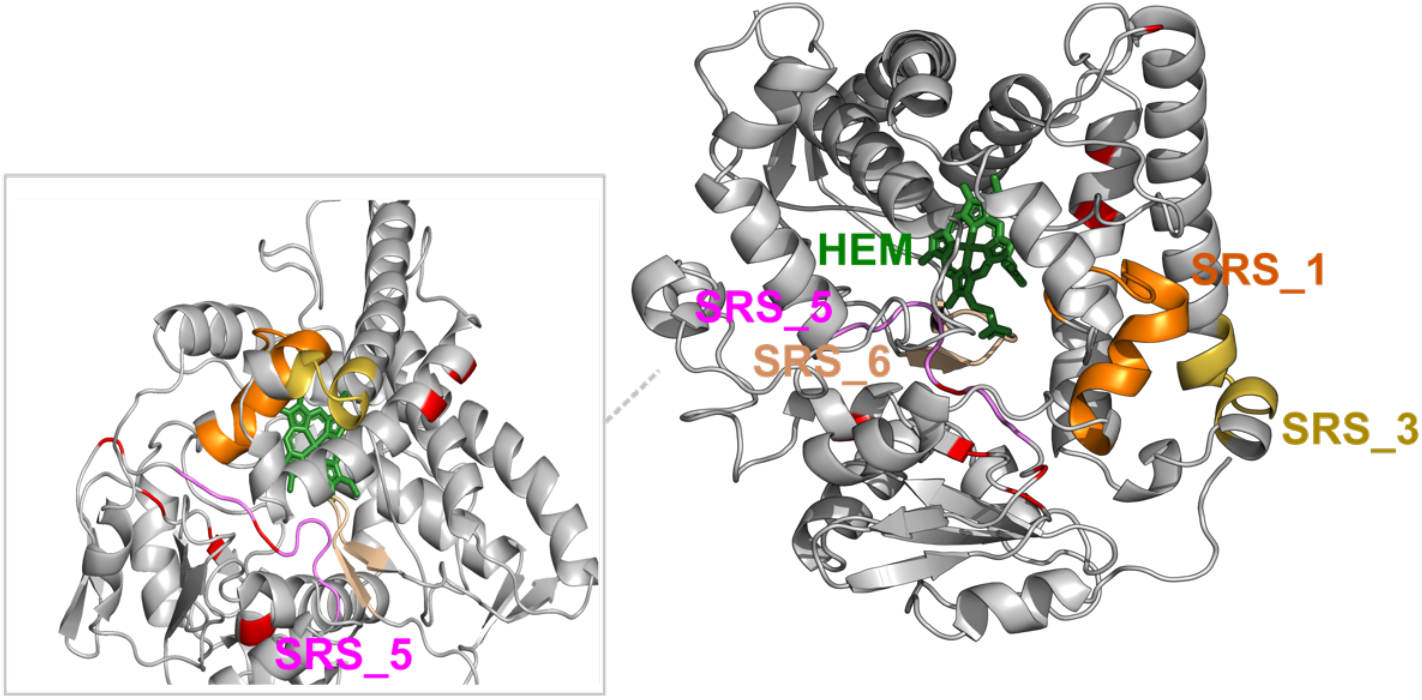
3D analysis of sites identified as being under positive selection in the avian CYP4B1. Positively selected sites identified by codeml site-models analyses (BEB, PP ≥ 0.95) are mapped onto a homology-based structural model of CYP4B1, shown in red. Residues overlapping previously defined substrate recognition sites (SRS) are hihglighted. SRS 1: orange, SRS 3: yellow, SRS 5: magenta and SRS 6: wheat.

#### CYP4F3

In CYP4F3, site-model analyses supported substantial heterogeneity in selective pressures among codon sites, with M3 fitting significantly better than M0 (Table S8). However, neither the M1a vs M2a nor the M8a vs M8 comparisons supported the presence of a positively selected site class, and ω estimates under M8 were constrained to one. Although the M7 vs M8 comparison was significant, this pattern is consistent with relaxation of selective constraint rather than adaptive evolution. No codons were identified with high posterior support under BEB. These results suggest that while selective constraints vary across CYP4F3, there is no clear evidence for site-specific positive selection.

#### CYP4F22

Analyses of CYP4F22 similarly revealed heterogeneity among codon sites, as indicated by the significant M0 vs M3 comparison (Table S9). In contrast, comparisons explicitly testing for positive selection (M1a vs M2a and M8a vs M8) were not significant, and ω estimates under M8 did not exceed one. Although the M7 vs M8 comparison was significant, this signal reflects variation in selective constraint rather than the presence of positively selected sites. A single codon (site 6) showed elevated posterior support under BEB (PP = 0.991 under M8), representing <0.3% of sites in the alignment (Table S10). This site does not overlap a known SRS, and PRIME analyses did not detect associated changes or conservation of amino-acid properties under the applied thresholds. Overall, evidence for positive selection in CYP4F22 is limited and localised.

#### CYP4V2

CYP4V2 showed clear and consistent evidence for site-specific positive selection. The M3 model provided a significantly better fit than M0, indicating substantial heterogeneity in selective pressures across sites (Table S11). Both M1a vs M2a and M7 vs M8 comparisons supported models allowing a positively selected site class, and this signal was confirmed by a significant M8a vs M8 comparison. Under M2a, approximately 4.7% of sites were inferred to evolve under positive selection (p_2_≈ 0.047), with ω_2_ estimates approaching 1.8. BEB analyses identified multiple codons with high posterior support (PP ≥ 0.95), including sites 157, 171, 243, 244, 246, 279, 349, 409, and 411 (Table S12). Several of these sites were located within predicted helices, and one site (246) overlapped substrate recognition site 3 (SRS3). PRIME analyses provided site-specific insight into the physicochemical properties associated with substitutions at these positions. Site 157, located within a predicted helix showed evidence for changes in amino acid composition, while bipolarity and secondary structure related properties were conserved. Site 171, also located within a helix, was identified as positively selected by BEB, but did not show evidence for significant changes or conservation of specific amino acid properties under the applied PRIME thresholds. Site 243, located within a helix, showed elevated posterior support under BEB but did not exhibit detectable property-level signals in PRIME. Site 244, also within a helix, similarly showed no significant PRIME-inferred property changes or conservation. Site 246, which overlaps substrate recognition site SRS3, showed conservation of amino acid composition, suggesting selective constraint on residue identity within this functional region. Site 279, not annotated to a specific secondary structure element, exhibited changes in residue volume, with no conserved properties detected. Site 349, located within a predicted helix, showed evidence for changes in bipolarity, amino acid composition, and residue volume, alongside strong conservation of secondary structure related properties. Site 409, located withis a predicted ?-strand, ehibited changes in amino acid composition and charge, with conservation of bipolarity. Site 411, lacking explicit secondary structure annotation, showed high posterior support under BEB but no associated PRIME-inferred property changes or conservation.

Taken together, thee results indicate that positive selection in CYP4V2 affects a defined subset of sites distributed across helices, a β-strand, and a substrate recognition region, with substitutions associated with heterogeneous and site-specific shifts in physicochemical proeprties. Given the presence of positively selected sites within SRS3 and multiple structured regions, the spatial distributions of these residues was examined using homology-based structural models to visualise their relative positioning within the protein (Fig 4).

**Fig 4.**
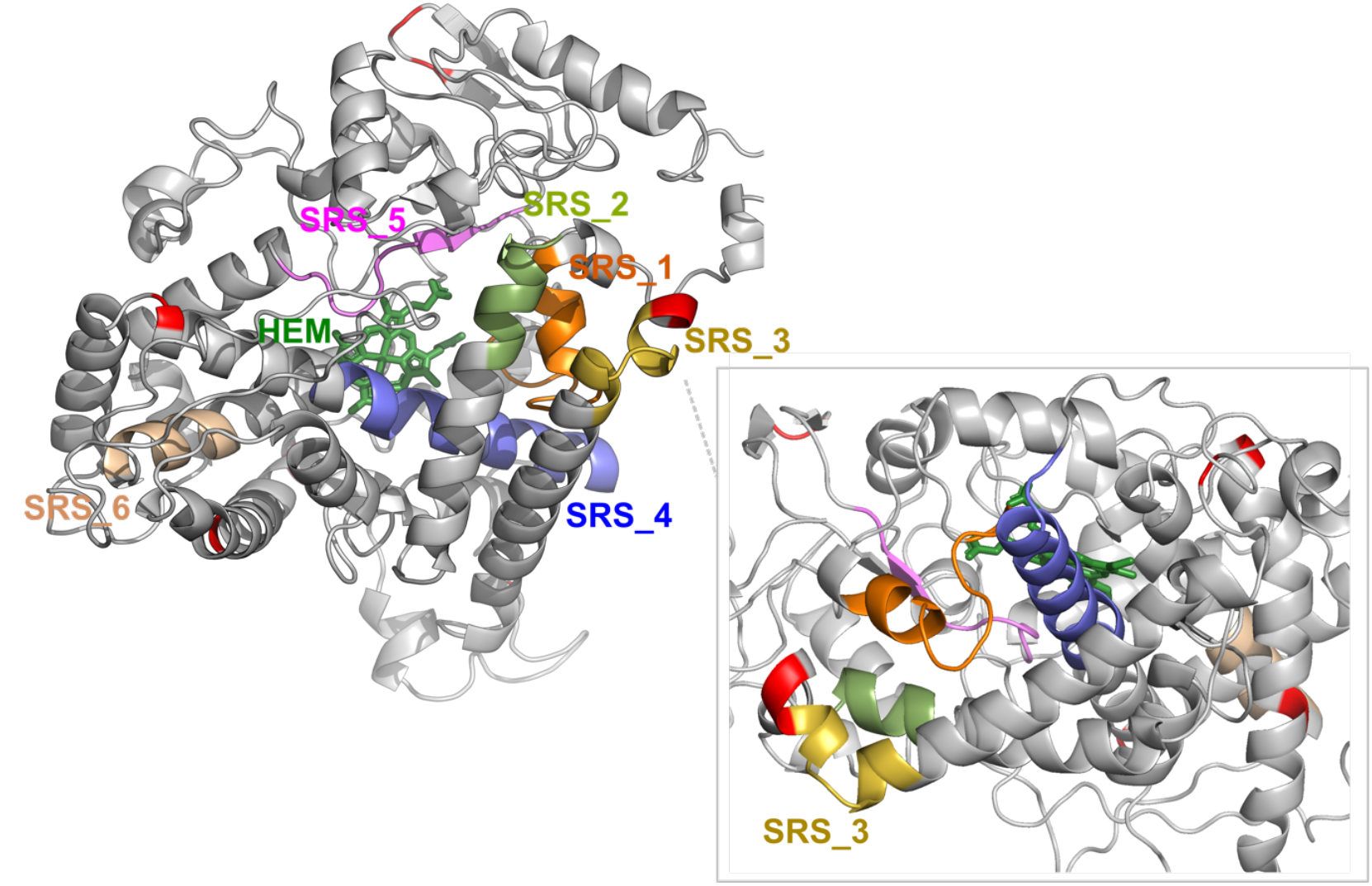
3D analysis of sites identified as being under positive selection in the avian CYP4V2. Positively selected sites identified by codeml site-models analyses (BEB, PP ≥ 0.95) are mapped onto a homology-based structural model of CYP4V2, shown in red. Residues overlapping previously defined substrate recognition sites (SRS) are hihglighted. SRS 1: orange, SRS 2: light green, SRS 3: yellow, SRS 4: blue, SRS 5: magenta and SRS 6: wheat.

### Trait-associated Variation in Selective Pressure Across Avian CYP4 Lineages

To examine whether selective pressures acting on CYP4 genes varied among ecological and life-history categories, branch models implemented in CODEML were applied using discrete trait assignments. For each gene, likelihood ratio tests (LRTs) compared a null model assuming a single ω across all branches with alternative models allowing trait-specific ω values. Traits examined included feeding habits, migratory behaviour, and primary lifestyle, as defined in the AVONET (Tobias et al. 2022) dataset. Significant LRTs were interpreted as evidence that selective pressures differ among trait categories, whereas non-significant results were taken to indicate broadly similar evolutionary constraints across lineages.

#### CYP4A4

Branch-model analyses of CYP4A4 provided limited evidence for trait-associated variation in selective pressure. No significant differences in ω were detected among feeding habit categories or primary lifestyle groups (Table S13). In contrast, the model allowing migration-specific ω values fit the data significantly better than the single-ω model. Under this model, migratory and partially migratory lineages showed lower ω values than non-migratory lineages, which exhibited the highest ω estimate. These results suggest that selective constraints acting on CYP4A4 may vary with migratory behaviour, although ω values across categories remained below one, indicating overall purifying selection.

#### CYP4A5

For CYP4A5, branch-model comparisons did not reveal significant differences in ω among feeding habit categories, migratory classes, or primary lifestyle groups (Table S14). Although alternative models estimated different ω values for some trait categories, none of the corresponding LRTs reached statistical significance. These results indicate that, despite clear evidence for site-specific positive selection in CYP4A5, lineage-level variation in selective pressure associated with the examined traits is limited.

#### CYP4B1

In CYP4B1, branch-model analyses identified significant variation in ω associated with feeding habits and migratory behaviour (Table S15). Models allowing separate ω values for carnivorous, herbivorous, and omnivorous lineages fit the data significantly better than the single-ω model, although estimated ω values across categories remained similar and below one. Migration-specific models also showed a highly significant improvement in fit, with modest differences in ω among migratory, partially migratory, and non-migratory lineages. In contrast, no significant differences were detected among primary lifestyle categories. Together, these results suggest that selective pressures acting on CYP4B1 vary among lineages associated with diet and migration, while remaining dominated by strong purifying selection overall.

#### CYP4F22

Branch-model analyses of CYP4F22 revealed limited evidence for trait-associated variation in selective pressure. No significant differences in ω were detected among feeding habit categories or primary lifestyle groups (Table S16). However, the migration-specific model provided a significantly better fit than the null model. Under this model, migratory lineages showed higher ω estimates relative to partially migratory and non-migratory lineages, although ω values remained below one across all categories. These results indicate that migratory behaviour may be associated with modest shifts in selective constraint acting on CYP4F22.

#### CYP4V2

For CYP4V2, branch-model analyses identified a significant effect of migratory behaviour on ω, whereas no significant differences were detected among feeding habits or primary lifestyle categories (Table S17). Under the migration-specific model, migratory and partially migratory lineages exhibited lower ω values than non-migratory lineages. Although all estimated ω values were below one, the significant LRT indicates that selective pressures acting on CYP4V2 differ among migratory classes.

Across avian CYP4 genes, patterns of molecular evolution differed among subfamilies. Evidence for site-specific positive selection supported by multiple site-model comparisons was detected in CYP4A5, CYP4B1, and CYP4V2, each of which harboured BEB-identified codons overlapping defined substrate recognition sites. In contrast, CYP4A4, CYP4F3, and CYP4F22 exhibited substantial heterogeneity in selective constraint across sites but lacked consistent evidence for codons evolving under positive selection. Branch-model analyses showed that lineage-level variation in ω was most frequently associated with migratory behaviour, whereas feeding habits and primary lifestyle displayed limited or gene-specific effects. Across all genes, migration-associated differences in ω remained below one and varied in their presence and magnitude among subfamilies, consistent with variation in the strength of purifying selection rather than lineage-specific positive selection.

## Discussion

The widespread retention of CYP4B1 across avian lineages contrasts with the repeated inactivation events documented for this gene in hominoids, indicating divergent evolutionary trajectories across vertebrate lineages. Experimental analyses in primates have identified several amino acid positions in CYP4B1 whose modification leads to reduced enzyme stability or catalytic activity, including residues located within the N-terminal core of the protein (Hüsken et al. 2025). Among these, Val71 occupies a conserved position within an N-terminal α-helix and contributes to hydrophobic packing that stabilises the overall fold of mammalian CYP4B1, with substitution at this site shown to markedly impair enzyme stability and function.

In the present analysis, a site homologous to Val71 was identified as being under positive selection in the avian CYP4A5 subfamily (Table S5 Site 58, located within α4 in Figure S2 PODCRI_CYP4A5), where it is occupied by methionine in several species. Structural alignment indicates that this residue lies within a conserved α-helical element, suggesting that it occupies a comparable structural position across CYP4 subfamilies. Although a Val-to-Met substitution represents a conservative change relative to Val-to-Gly substitution described in Denisovans, the detection of positive selection at this position in CYP4A5 indicates that even subtle alterations at conserved structural sites may be subject to adaptive fine-tuning. CYP4A enzymes are primarily associated with fatty-acid and lipid metabolism, and modest changes affecting protein stability or efficiency could provide a mechanism for adjusting enzymatic performance while preserving overall catalytic architecture. The identification of positive selection at a structurally conserved site shown to be functionally important in another CYP4 subfamily therefore supports a broader model in which adaptive evolution within the CYP4 family can act on shared structural “lever points”, with subfamily-specific selective pressures determining where and how such positions are targeted.

CYP4V2 also showed broad retention across avian species. Studies in humans have shown that disruption of this gene has substantial physiological consequences, as loss-of-function mutations in CYP4V2 cause Bietti crystalline corneoretinal dystrophy, a rare inherited retinal degeneration characterised by abnormal lipid accumulation (Li et al. 2004). In humans, CYP4V2 catalyses ω-hydroxylation of medium- and long-chain fatty acids, including polyunsaturated substrates, and is expressed in the retina as well as multiple other tissues. The broad retention of CYP4V2 across birds therefore suggests that this subfamily may contribute to metabolic processes in avian species. CYP4F3 and CYP4F22 showed little evidence for site-specific positive selection, suggesting more stable evolutionary dynamics across this subfamily.

Our analyses indicate that selective pressures acting on avian CYP4 genes are most evident at the level of individual codon sites, with more limited modulation of selective constraint across lineages. Similar distributions of purifying selection and localised functional diversification have been reported for cytochrome P450 proteins (Gotoh 1992; Montellano 2015).

Evidence for positive selection was detected in a subset of CYP4 genes and involved only a small proportion of codon sites. The localisation of these sites to substrate recognition regions and structured elements is notable, as substrate recognition sites have long been identified as focal points of functional diversification in cytochrome P450 enzymes (Gotoh 1992). Comparative structural studies have shown that cytochrome P450 enzymes share a broadly conserved overall fold, with sequence and structural differences often observed in regions associated with substrate binding (Hasemann et al. 1995; Thomas 2007).

Together, these observations indicate that sequence variation associated with functional diversification in CYP4 genes is concentrated at residues involved in substrate interaction, within a broadly conserved protein fold.

Structural mapping based on homology models indicated that positively selected residues are embedded within defined secondary structural elements and, in several cases, within canonical substrate recognition sites. Although homology models alone cannot fully resolve functional mechanisms, the spatial distribution of selected sites is consistent with models proposed for other cytochrome P450 families, in which adaptive change involves fine-scale adjustment of residue properties within a conserved structural framework (Hasemann et al. 1995; Almeida et al. 2016). Analyses of amino-acid properties indicated that substitutions at these sites are associated with specific changes or conservation of physicochemical properties, supporting constrained optimisation rather than random amino acid replacement.

At the lineage level, branch-model analyses revealed modest variation in selective pressure. Migration was the trait most consistently associated with differences in ω, but these differences remained below one across all genes, indicating modulation of purifying selection rather than episodic or lineage-specific positive selection.

Comparable associations between life-history or ecological variables and background evolutionary rates have been reported in large-scale comparative analyses across mammals, suggesting that trait-linked shifts in selective constraint can arise without invoking lineage-specific adaptive evolution (Nikolaev et al. 2007; Bromham 2009). Lineage-level variation in selective pressure and site-specific signatures of positive selection showed little overlap across genes.

Overall, these results suggest that the evolution of avian CYP4 genes is shaped by strong background purifying selection, with adaptive diversification occurring through localised, site-specific changes affecting a limited number of residues within functionally relevant regions, while ecological variation among lineages primarily influences the strength of selective constraint rather than driving lineage-specific adaptation.

## Materials and Methods

### Whole-genome identification of CYP4 Gene Sequences

To characterize avian CYP4 genomic diversity, we conducted tBLASTn searches on 363 avian genomes from the Bird 10,000 Genomes (B10K) Project dataset using as query CYP4 subfamily protein sequences annotated in Ensembl for chicken (*Gallus gallus*), turkey (*Meleagris gallopavo*), anole lizard (*Anolis carolinensis*), frog (*Xenopus tropicalis*), zebrafish (*Danio rerio*) and human (*Homo sapiens*). From the CYP4 sequences retrieved, psudogenes were removed and only nucleotide sequences with more than 1125bp and high identity (e-value <1e^-5^) were used for further analysis.

### CYP4 Gene Subfamily Data Sets and Phylogenetic Analysis

Codon-based alignment was constructed of all identified avian CYP4 nucleotide sequences in MAFFT (Katoh and Standley 2013), along with some reference sequences from public databases (Ensembl release 112: https://www.ensembl.org and NCBI: https://www.ncbi.nlm.nih.gov, last accessed May 2024). These references included representative CYP4 subfamilies from human (*Homo sapiens*), anole lizard (*Anolis carolinensis*), chicken (*Gallus gallus*), Chinese medaka (*Oryzias sinensis*), Chinese softshell turtle (*Pelodiscus sinensis*), Eastern brown snake (*Pseudonaja textilis*), tropical clawed frog (*Xenopus tropicalis*), medium ground finch (*Geospiza fortis*), and spotted gar (*Lepisosteus oculatus*). A total of 1056 avian sequences were aligned along with 42 CYP nucleotide sequences from reference species followed by a construction of Maximum Liklihood (ML) phylogeny. This ML phylogeny assumed a General Time Reversible (GTR) evolutionary model, with a proportion of invariable sites (GTR+I+G4), as determined by ModelTest-NG (Darriba et al. 2019). The ML phylogeny was estimated using IQ-TREE 2 (v2.3.6) (Hoang et al. 2018; Minh et al. 2020) with 1000 bootstrap replicates.

CYP4 nucleotide sequences from each well-defined clade in this ML phylogeny were retrieved and grouped as CYP4 subfamily datasets. Databese reference sequences were removed from the CYP4 subfamily datasets and profile HMMs were generated from reference CYP4 subfamily sequences using HMMER (Finn et al. 2011) and each of these CYP4 subfamily sets were then aligned against these profile alignments using MAFFT. Several MAFFT alignements were constructed, that led to the identification of short incomplete sequences, which were removed from the respective datasets. Each CYP4 subfamily datasets was checked closely to ensure only one CYP4 sequence per avian species was present in each set. These were further tested for recombination using RDP4 (Martin et al. 2015) with default settings and seven algorithms (BootScan, Chimaera, GENECONV, MaxChi, RDP, SiScan, and 3Seq). CYP4 sequences with evidence of gene conversion events or recombination (*p*-values < 0.05) were removed. Fifteen CYP4 subfamily datasets were identified following this approach: CYP4A4, CYP4A5, CYP4A6, CYP4A10, CYP4A11, CYP4A22, CYP4A25, CYP4B1, CYP4B7, CYP4F3, CYP4F4, CYP4F8, CYP4F11, CYP4F22 and CYP4V2. The details of which species is represented in each dataset can be found in Supplementary File 1 (sheet: Avian_CYP4_subfamily_dataset).

### Ancestral Reconstruction Analysis

To better understand the evolutionary process of avian CYP4 subfamilies, ancestral sequence reconstruction was performed with the COUNT software (Csűös 2010) using CYP4 subfamilies identified above and the species tree from B10K Project (https://cgl.gi.ucsc.edu/data/cactus/363-avian-2020-phast.nh) (Feng et al. 2020). The CYP4 subfamily genes present in each avian species were converted into binary numerical gene profiles (0 for absent, 1 for present) and the data was then analyzed using the Dollo parsimony model (Farris 1977) with default parameters.

### Selection Analysis of Avian CYP4 Subfamilies

Analyses of selective pressures acting on CYP4 gene sequences were performed using the CODEML program implemented in PAML (v4.10.7), which estimates the ratio of nonsynonymous to synonymous substitution rates (ω = *dN/dS*) under maximum likelihood (Yang 2007; Bielawski et al. 2016). This allows testing whether particular codon sites or evolutionary lineages deviate from neutral expectations, providing evidence for positive selection or heterogeneous selective pressures across the phylogeny. Site-models, branch-models, and their corresponding null models were fitted following standard procedures to evaluate variation in ω among codons and lineages. Following the recommendations in (Jeffares et al. 2014), only CYP4 subfamily datasets with more than 10 species (sequences) were used for the analyses. Six CYP4 subfamily datasets (CYP4A4, CYP4A5, CYP4B1, CYP4F3, CYP4F22 and CYP4V2) fulfilled this criteria. For every CYP4 subfamily dataset, the codon-based alignment and the corresponding unrooted subset tree generated from the B10K species tree (363-avian-2020-phast.nh) were submitted to the site-model analysis.

For site-model analyses, we compared nested models that differ in whether they allow classes of codons with ω > 1. Specifically, null and alternative models were contrasted using likelihood ratio tests (LRTs): M0 vs. M3, M1a vs. M2a, M7 vs. M8, and M8a vs. M8. Whenever the LRT was significant (*p*-value < 0.05), codon sites with Bayes Empirical Bayes posterior probabilities > 0.95 under models M2a or M8 were considered to be under positive selection. CYP4 subfamilies showing evidence of sites under positive selection in the site-model analysis were subsequently analyzed using branch-models to assess whether selective pressures varied among major avian lineages. To account for uncertainity in the evolutionary history of ecological traits, we reconstructed trait evolution on the B10K avian species tree by stochastic character mapping (SCM). SCM samples complete histories of character transitions along branches under an explicit continuous-time Markov model, thereby incorporating both uncertainity in ancestral reconstructions and variation in the timing and frequency of trait changes. This approach allows trait states to vary along branches rather than being restricted to nodes, which is particularly relevant for traits that may have undergone multiple transitions within lineages (Pagel 1994; Huelsenbeck et al. 2003). The SCMs were generated using the R packages phytools and geiger, following the method of Bollback (Bollback 2006; Revell 2012; Pennell et al. 2014). The unrooted B10K species tree was labeled according to the mapping results and was trimmed to contain only the species present in each CYP4 subfamily dataset. In these analyses, the branch-model configuration allowed separate ω values to be fitted for foreground branches representing distinct avian groups, and model fit was evaluated by comparing models that allow different ω ratios among branches to the one-ratio null model. Across all analyses (for both site-models and branch-models), the F3×4 codon frequency model was applied. To reduce the risk of convergence on local likelihood optima, each model was run multiple times with different initial values of *k* (transition/transversion rate ratio) and ω. Parameter estimates and LRT statistics were taken from the replicate with the highest log-likelihood score. The avian traits of each species (life style, migratory status and feeding habbits) are based on the classification in AVONET (Tobias et al. 2022). Details of avian traits can be found in Supplementary File 1 (sheet: Avian_traits).

The Property-informed Models of Evolution (PRIME) method implemented in HyPhy(v2.5.85) (Pond et al. 2005) was used to assess site-specific physiochemical and amino-acid properties that are preserved or altered during evolution. PRIME models the evolution of coding sequences on a fixed phylogeny by integrating site-specific substitution rates with quantitative descriptors of amino-acid properties, including polarity (bipolarity), charge, residue volume, secondary-structure propensity, and compositional class (Atchley et al. 2005; Conant et al. 2007). For each site, the method evaluates whether observed substitutions across the phylogeny show a consistent tendency to increase or decrease a given property, relative to expectations under a neutral model that assumes no property-specific directional bias. Evidence for property-based selection is assessed using likelihood ratio tests comparing property-informed and property-independent models, and statistical significance is evaluated using both site-level *p*-values and false-discovery-rate (FDR) correction. It reports the set of properties showing significant directional change, the estimated strength and direction of selection on each proeprty, and properties inferred to be conserved. From the sites reported by PRIME, we only considered and analyzed sites that were classified as being under positive selection in the CODEML site-model analysis.

Cytochrome P450 enzymes comprise a large and diverse gene superfamily, yet share a conserved three-dimensional fold that accomodates substrate binding and catalysis across a wide range of chemical substrates. Within this conserved architecture, substrate recognition sites (SRS) are regions close to ligands in the tridimensional (3D) structure of the protein and have been shown to influence ligand binding and contribute to differences in substrate specificity and functional diversification among CYP enzymes (Hasemann et al. 1995; Johnson and Stout 2013). CYP4 SRS were defined based on established cytochrome P450 annotations inferred from comparative sequence analysis (Gotoh 1992) and through large-scale structure-substrate mapping (Zawaira et al. 2011). Following this approach, six SRS were mapped on the CYP4 amino acid sequences (Supplementary Table S1 and Supplementary Figure S2). Numerous cytochrome P450 crystal structures have been determined across taxa, with several hundred entries currently available in the Protein Data Bank (RCSB PDB: http://www.rcsb.org/) (Berman et al. 2000). However, structural representation remains uneven across CYP families, and rabbit CYP4B1 (Hsu et al. 2017) represents the only experimentally determined X-ray crystal structure from the CYP4 family to date.

To gain understanding on the spatial organization and structural relationship, we performed homology modeling of the 3D structure of avian CYP4, using the rabbit CYP4B1 structure from PDB (PDB code: 5T6Q) as template in MODELLER (v10.8) (Webb and Sali 2016). Substrate recognition sites mapped to the CYP4 sequences and positively selected sites identified for each CYP4 subfamily dataset in the CODEML site-model analyses were mapped onto the 3D structure. Visualization and superimposition of the 3D structures was done with PyMOL (v3.1.0) (http://www.pymol.org).

## Supplementary Material

Supplementary_File_1.xlsx

Supplementary_Fig_Tables.pdf

## Acknowledgements

This research was supported by The National Centre for the Replacement Refinement & Reduction of Animals in Research(NC3Rs) CRACK IT project DARTpaths [project number CRACK-ITDP-P1-2].

